# Perfect Counterfactuals for Epidemic Simulations

**DOI:** 10.1101/451153

**Authors:** Joshua Kaminsky, Lindsay T. Keegan, C. Jessica E. Metcalf, Justin Lessler

## Abstract

Simulation studies are often used to predict the expected impact of control measures in infectious disease outbreaks. Typically, two independent sets of simulations are conducted, one with the intervetnion, and one without, and epidemic sizes (or some related metric) are compared to estimate the effect of the intervention. Since it is possible that controlled epidemics are larger than uncontrolled ones if there is substantial stochastic variation between epidemics, uncertainty intervals from this approach can include a negative effect even for an effective intervention. To more precisely estimate the number of cases an intervention will prevent within a single epidemic, here we develop a ‘single world’ approach to matching simulations of controlled epidemics to their exact uncontrolled counterfactual. Our method borrows concepts from percolation approaches prune out possible epidemic histories and create potential epidemic graph that can be ‘realized’ to create perfectly matched controlled and uncontrolled epidemics. We present an implementation of this method for a common class of compartmental models, and its application in a simple SIR model. Results illustrate how, at the cost of some computation time, this method substantially narrows confidence intervals and avoids non-sensical inferences.

## 1 Introduction

Dynamic models are frequently used to assess the likely impact of disease control strategies. These exercises range from modeling the impact of a new intervention or strategy on an established pathogen [1, 2, 3], to models of the containment and control of emergent epidemics [4, 5, 6]. While deterministic models are frequently used [7, 2, 8], stochastic simulations are increasingly common as they can account for both uncertainty in the underlying parameters and the random nature of the disease process [6, 9, 5, 10]. In both stochastic and deterministic models, the impact of interventions are typically determined by comparison of simulations with and without the intervention.

In the deterministic setting, this comparison is straight forward, as with a given set of parameters and starting conditions the epidemic will always behave exactly the same; hence, any comparison between an intervention scenario and its non-intervention ‘counterfactual’ can only be attributed to the intervention itself. When stochastic models are used, things become more complicated. Typically, two sets of simulations are conducted, one with the intervention and one without, and then the distribution of outcomes from the two sets are compared to estimate the intervention effect. Because these are independent sets of simulations, there may be some simulations in the non-intervention scenario where the disease dies off quickly due to random chance, and fewer cases occur than the majority of intervention simulations. Likewise, there may be cases in the intervention scenario where large numbers are infected through sheer ‘bad luck’ for the virtual populations involved. If these stochastic effects are large, they may lead uncertainty intervals in effect estimates to include the intervention having no effect, or even a negative impact, even if the intervention is modeled in such a way that it can only have a beneficial effect. For example, the results of a study of measles vaccination strategies by Lessler et al. appears to leave open the possibility that more cases of measles could occur in a country if supplementary vaccination activities were conducted than if those campaigns had not occurred [3]. Likewise, Rivers et al. is consistent with low coverage of pharmaceuticals causing additional cases of Ebola when compared to no coverage, even though the model assumes they have a beneficial effect [11]. These effects will be exacerbated if the processes being modeled are complex or we are simultaneously sampling over parameter uncertainty.

The results of this independent simulation approach have a very specific interpretation: they represent the difference between what we expect to be observed in an uncontrolled epidemic compared to a completely independent epidemic where the intervention occurred (conditional on the starting conditions). However, what we often want to know is what would have happen had the intervention occurred in the exact same epidemic. This is analagous to the problem of counter-factual inference in randomized trials and observational studies, where we take one set of individuals (or populations) as a stand in for what would have happened in the counterfactual situation that they had (or had not) experienced some exposure. A number of techniques of trial design and statistical analysis have been developed to help real world studies better approximate the true counterfactual situation ([12, 13, 14] are just a few examples of the large literature on the subject). However, in computational simulations it is possible to take a more exact approach.

Here, we present a method for simulation of direct counterfactuals to stochastic simulations using principles borrowed from the percolation approach to epidemic simulation [15]. We illustrate this ‘single-world’ approach using simulations of interven-tions against an influenza like illness, and outline how the general approach can be adapted to a wide variety of disease systems and simulation methodologies.

## 2 Methods

### Overall Approach

Our goal is to simulate an uncontrolled epidemic and then simulate one or more controlled epidemics that are 100% consistent with the uncontrolled epidemic. That is, all events (e.g., transmissions, recoveries, etc.) that occur in the uncontrolled epidemic also occur in its controlled counterpart unless precluded (directly or indirectly) by the intervention; and no stochastic events that did not occur but were possible in the uncontrolled epidemic happen in its controlled counterpart unless explicitly caused by an intervention. To accomplish this, we take an approach that first prunes events (Figure 1) according to a model based stochastic simulation to create a ‘potential epidemic graph’ (PEG) (Figure 2), and then use this graph to ‘realize’ consistent epidemics in controlled and uncontrolled scenarios (Figure 3).

**Figure 1:**
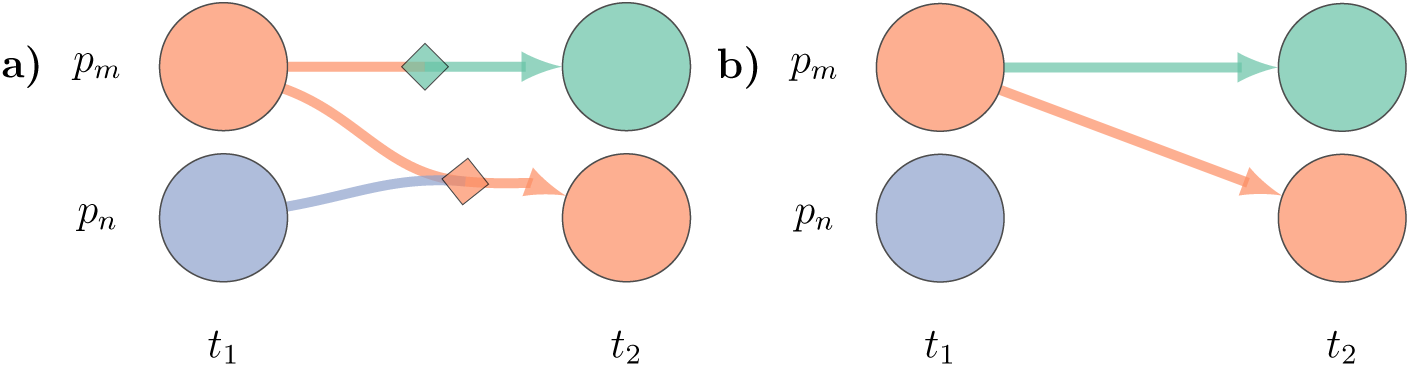
*Anatomy of an event*. Each event has pre-conditions based on the state of one or more individuals *p* in the population at time *t*, a probability of occurrence, and outcome that sets the state of an individual at time *t*_1_. (a) In its full representation an event is represented as a node on the graph (diamonds) with incoming edges representing the preconditions individuals (circle nodes) must meet for that event to occur. In this case both events require individual *p*_*m*_ to be *infectious* (orange) and the bottom event requires individual *p*_*n*_ to be *susceptible* (purple). The outgoing edge captures the outcome of the event, in this case *removed* (green) and *infectious*. (b) We use a reduced representation where the event nodes are implicit. Edges are colored by their outcome, and infectious edges (orange) carry an implicit precondition that the target is susceptible. Here, the reduced representation in panel (b) is equivalent to the full graph in panel (a).

**Figure 2:**
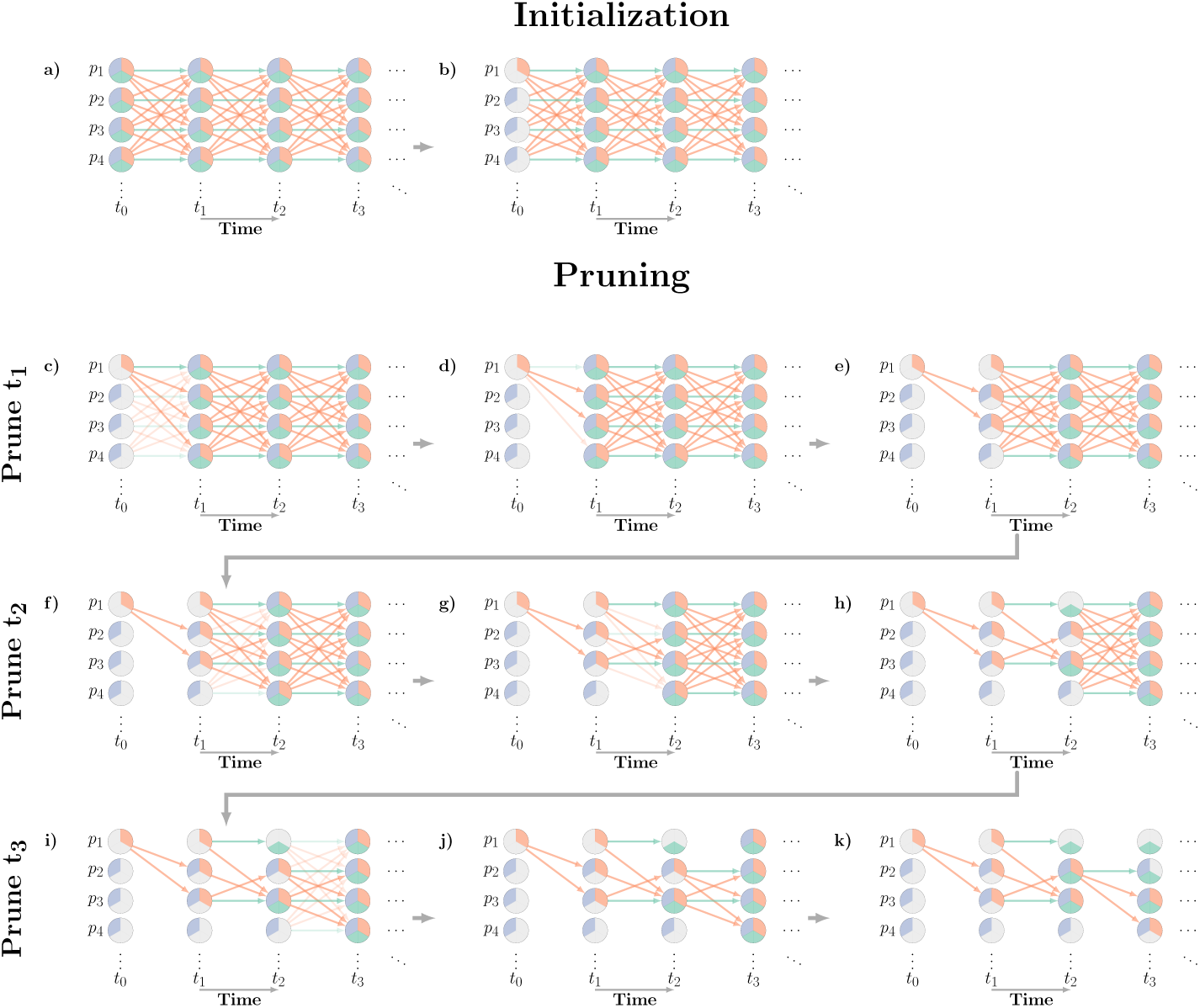
*‘Single-world’ simulation process*. To simulate an initial epidemic and interventions, we start with a (implicit) complete graph. We represent potential states using colored wedges; an individual can be *susceptible* (purple pie slice), *infectious* (orange) or *removed* (green). We start by using our initial conditions to eliminating all but one potential state for each individual in the population a state at time *t*_0_ (b), here infecting person *p*_1_. We next prune all events with at least one precondition which we know is not satisfied by the initial state (c). We then prune those events stochastically selected not to occur by our underlying infection model (d). We set the possible states of each individual at *t*_1_ based on the remaining events in the graph, so each person’s set of potential states encompasses both the outcomes of any remaining events, and outcomes of their absence (e). We now repeat steps c-e for events between *t*_0_ and *t*_1_ (f-h), *t*_1_ and *t*_2_ times 2 and 3 (i-k) and so on. Note that when we prune the events with unattainable preconditions, (i) we still keep the those for *p*_1_ and *p*_2_ infecting *p*_3_, even though *p*_3_ was potentially *infectious* at *t*_1_, because *susceptible* is still a possible state for *p*_3_ at *t*_2_ in an intervention scenario (i.e., removing the transmission event at *t*_1_ could lead to a scenario where *p*_3_ is susceptible at *t*_2_.) This final potential epidemic graph (PEG) can then be used to obtain simulated epidemics with and without interventions (see Figure 3).

**Figure 3:**
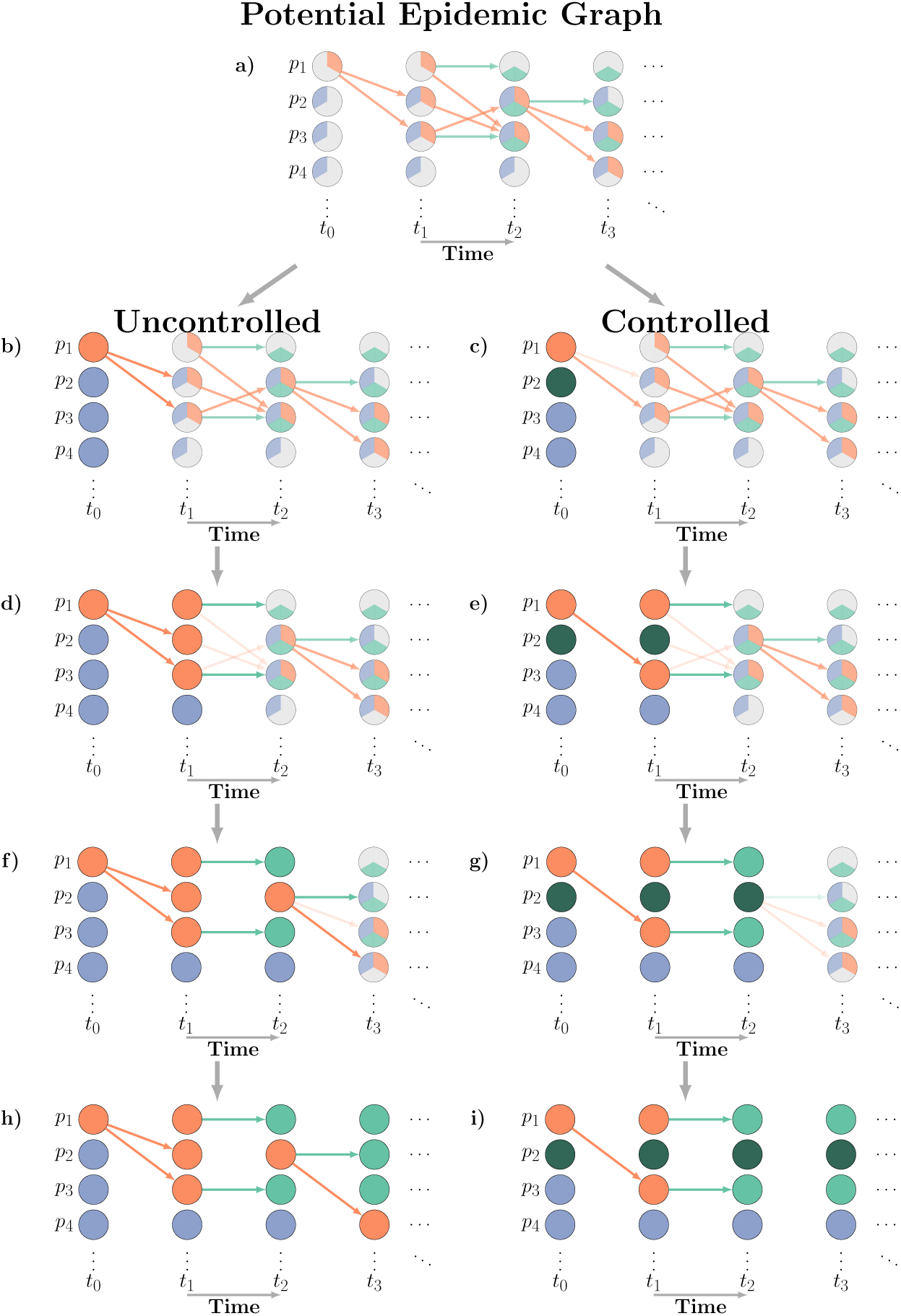
*‘Single-world’ simulation process (continued)*. In order to measure the impact of the intervention on the epidemic, we use the potential epidemic graph to construct two epidemics: one uncontrolled (left) one with intervention (right). We start by setting the actual state (denoted by coloring the whole circle) of each person at time *t*_0_ to their initial condition, and then allowing the change their initial states, in this case setting *p*_2_ as *vaccinated* (dark green). We then prune events between *t*_0_ and *t*_1_: removing inconsistent events and allowing the intervention to stochastically remove events (b,c). We then use those events to determine the actual state of each person at *t*_1_, allow the intervention to alter that state, then prune inconsistent events again (d,e). We repeat the process for times *t*_2_ (f,g), *t*_3_ (h,i), and so on. This final graph can be used to extract our outcome of interest. Any graphs made from the same PREPresent the results of different interventions in a ‘single-world’.

Specifically, our approach begins with an implicit complete graph that includes a representation of all possible events (Figure 2). In the complete graph, there are two types of nodes: one representing individuals at times, and one representing events (Figure 1). A node representing an individual at a particular time is considered to potentially belong to any of a set of *potential states* (e.g., susceptible, infectious, removed). Directed edges represent dependence. An edge from an individual node to an event node means that individual node has to be in a particular state (or set of states) as a *precondition* for that event. An edge from an event node to an individual node means that an *outcome* of that event is a change in the state of the target node (e.g., immune is the outcome of a recovery event). The underlying epidemic model assigns each event a *probability* of occurring if its preconditions are met. In the case where all events represent either a transition in an individual’s state dependent only on their prior state (e.g., recovery events) or infections resulting from contact between individuals, a simplified representation is possible (Figure 1). In this simplification, events are represented by edges and infection events have the implicit precondition of the target being susceptible at the previous time point. For clarity in text and figures, we will focus on this simplified representation for the rest of this manuscript, but it should be noted that the full representation allows for a greater diversity of models.

As illustrated in Figure 2, to create the PEG, we first assign individuals an initial state at time 0 (equivalent to removing all but one of their potential states). We then prune from the complete graph all of the events whose preconditions are inconsistent with the remaining potential states of each individual at time 0. Next, we stochastically prune the remaining events based on their probability. We then remove all states at time 1 that are inconsistent with both the states at time 0 and the remaining events. This process is then repeated iteratively until a predefined time limit is reached.

Once the PEG is created, an uncontrolled epidemic can be realized by iterating through times assuming all events whose preconditions are met occur and setting each event’s target node to the outcome of that event (Figure 3). To simulate an intervention, the same process is performed, but events are removed or node states changed probabilistically based on the intervention process.

Below, we describe how this process can be performed for disease systems specified by certain compartmen-tal models.

### Compartmental Models to PEGs

One of the most common ways to specify a epidemic model is by as system of probabilistic state (i.e., compartment) transitions, such as a stochastic SIR model. These models most commonly include two types of transitions: *contact transitions* triggered by infectious contacts and *independent transitions* resulting from the natural history of infection (e.g., recovery) or demographic changes (e.g., death) (Figure 4).

**Figure 4:**
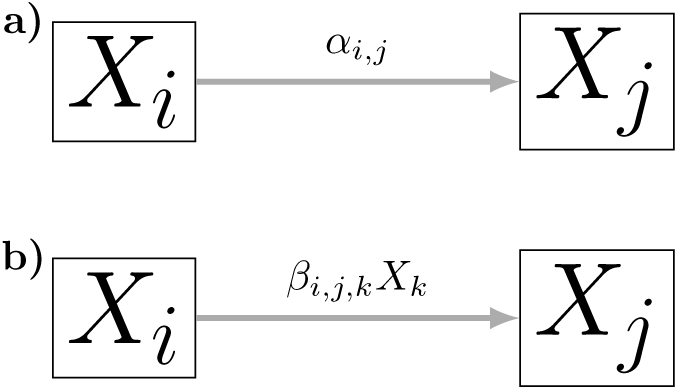
Two types of transitions commonly present in compartmental models: a) independent transitions occur with probability *α*_*i,j*_ independent of the state of the model, and **b**) contact transitions occur between two specific people with probability *β*_*i,j,k*_.

Here, we consider the set of models with *K* states whose behaviour can be described using only independent and contact transitions (this includes most SIR type models), and demonstrate how to construct the complete graph from these types of events. We assume that the probability of independent transitions between state *i* and *j* is *α*_*ij*_, and that the probability of contact transitions between *i* and *j* dependent on the number of people in state *k* is *β*_*i,j,k*_. In most models, *α*_*ij*_ will be 0 for the majority combinations of *i* and *j*, and, likewise, *β*_*i,j,k*_ will be 0 for the vast majority of combinations of *i, j*, and *k*.

To construct the (implicit) complete graph we create a node for each individual at each time point and assign their set of potential states to include all *K* states. We then add events between nodes for all pairs of adjacent time points. For each type of independent transition, we create an event between every individual and themself in the next time step. Similarly, for each contact transition, we create an event between each individual and every other individual in the next time step. To set the initial conditions, for each compartment *i* we set *X*_*i*_ individual nodes to have *i* as their only possible state at time 0. However, in some cases we may want to allow additional states if considering an intervention that can assign those states. This complete graph can then be used to construct the PEG using the algorithm described above.

### Interventions

Interventions can remove events (e.g., eliminate infection events) or change the states of individuals (e.g., make someone immune by vaccination). As outlined above, interventions are applied as we iterate over times when re-alizing the PEG, but for this process to work the PEG must also subsume the courses of the epidemic that are possible under the intervention (i.e., edges pruned as impossible never occur in the epidemic).

Therefore, in order to eliminate potential states when pruning, we need to make some assumptions about potential control strategies. Otherwise, the control strategy could change any node to any state at any time. Hence, the algorithm works best if we restrict the set of states control measures can cause an individual to enter, *W*, and the types of events control measures can remove, *Z*. These assumptions aid pruning by guaranteeing no individual can enter a state not in *W* unless caused by an event, and that outcomes caused by events in *Z* will always occur if their preconditions are met. The latter allows us to prune states that must be exited, from the set of possible states (e.g., if recovery is in *Z* and recovery occurs on individual *i* at time *t*, then there is no way *i* can be infectious at time *t* + 1 unless an infection event occurs or infection is in *W*).

### Simple Example

We demonstrate our proposed method using a simple *susceptible* (S), *infectious* (I), *removed* (R) compartmental model in a closed population, with a force of infection *βSI* and a recovery rate of *γ*. We set the parameters of the model for a disease similar to influenza 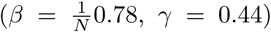 [16]. We consider the impact of three different control strategies, selected to capture common ways in which interventions work: antivirals, which reduce the period of infectiousness; hand washing, which reduces the probabil-ity of transmission; and vaccination, which removes people from the susceptible pool.

We ran 1,000 simulations using both classical techniques and the single-world approach for each intervention, on a population of 4, 000 for 100 days with 5 initial infected and calculated the estimated number of cases averted. To model antivirals, we assume that 10% of people will take antivirals if they are infected reducing their average duration of infection by 0.7 days [17], and therefore change the state of those individuals to *removed* if they are *infectious* with probability 20.1%. To model hand-washing, we assumed a 5% reduction in transmissibility, and therefore modeled it by removing 5% of transmission events. We assumed that 10% of people are given vaccine at the start of the epidemic, and that 33% of those are successfully vaccinated and start the epidemic in a *vaccinated* state.

We have posted a software package, `cfepi`, on `CRAN`, which implements our method. It follows the procedures outlined above assuming that interventions never put anyone in a non-terminal state and that interventions do not prune independent events. Time calculations for our method were made using this package.

## 3 Results

The time complexity of the single world approach is *O*(*TN*^2^) for both the construction of the PEG and realization, where *T* is the number of times steps and *N* is the number of nodes (i.e., it is limited by the number of possible combinations of individual pairs at each time-point). In practice, realization is far faster due to the extensive pruning during construction of the PEG. Storing the PEG can also take significant space, depending on the model and amount of pruning being done. In comparison, a standard implementation of a stochastic SIR model based on binomial draws is *O*(*T*). In our implementation, the example SIR disease system in a population of 400, 000 requires just less than a minute to create one PEG, < 20 seconds for each realization, and ~ 2GB of storage (significant speedups may be possible using GPU based parallelliza-tion). This is compared to < 1 second to run an iteration a standard stochastic SIR model. However, there is some reduction in the number of simulations needed due to the reduced variance in the intervention effects in the single-world approach (a reduction of about 2.5 times in this example).

Though it comes at a computational cost, the benefits for estimating intervention effects using the single world approach is evident in our simulations. For all of our illustrative interventions, we see virtually the same point estimate of impact in the single-world approach and standard approaches, but the single world approach gives confidence intervals that are both narrower and do not include negative effects. In a population of 4,000 (average uncontrolled epidemic size 2,849), the single-world approach estimated antivirals to prevent an average of 121 (90 % CI: 3–208) cases, versus 99 (90 % CI:-122–318) in the standard approach. The single-world approach estimated hand washing to prevent an average of 172 (90 % CI: 35–272) cases, versus 179 (90 % CI: −47–374). The single-world approach estimated vaccination to prevent an average of 196 (90 % CI: 79–287) cases, versus 215 (90 % CI: −8–412). Differences between single-world and indepdent approaches are even more pronounced if we compare intervention effects at different points during the epidemics (Figure 5)

**Figure 5:**
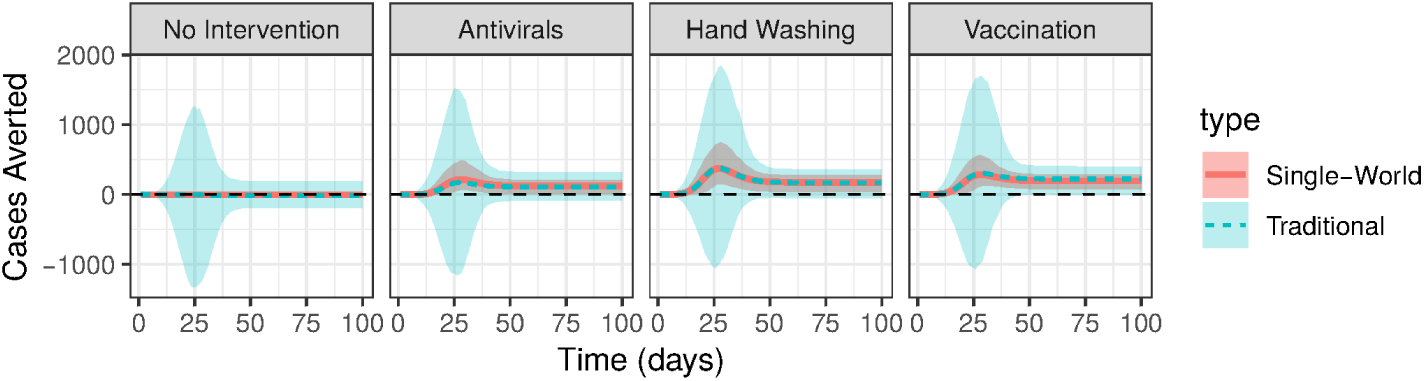
Time series showing the number of cases averted at each time caused by the intervention calculated using our method (single-world) and a standard method. Shaded regions denote 90% confidence intervals.

## 4 Discussion

In this paper, we present a method that allows simulation studies to more exactly estimate the precise impact of an intervention by comparing epidemic simulations with their exact ‘counter-factual’. We demonstrate the ability of our method to factor away cross-epidemic variation, which can lead to counterintuitive results, such as uncertainty intervals that suggest that an intervention programmed into the model to only have a beneficial effect has some small chance of having a negative one. We further show our method is tractable when applied to many familiar models and control strategies, and have implemented software to aid in its use in common cases.

This ‘single world’ approach of matching controlled epidemic simulations with their exact uncontrolled counterfactual answers a subtly dif-ferent question than comparisons between independently simulated sets of epidemics with and without control. The single world method estimates *how many cases we expect to be prevented by using a control measure;* while the independent simulation approach estimates *the expected difference in size between two different epidemics, one with and one without the control measure*. To put it in more concrete terms, an example of the first type of question is, “How many cases of influenza will we prevent next year if we increase vaccination rates by 15%?”, while the an example of the latter is, “How many fewer influenza cases do we expect next year compared to last if vaccination increases by 15%?”. Both are important public health questions, but authors are not always clear which they are answering, with a tendency to write as if they are answering the former, while performing simulations that answer the latter. This may seem unimportant, as in their expectation they have the same answer (ignoring season-to-season inter-dependence in influenza epidemics for the sake of argument); but, as illustrated by our simple examples, their overall distributions can differ substantially, and in ways that might lead one to discard an intervention that has an important effect. While in our simple SIR model the intervention effect had to be small for the predicted effects to cross null, in more complex systems this can happen even for interventions with a large impact (as in [3]).

We are not the first group to examine the refined counterfactual question. Kenah and Miller use percolation to measure the effects of a single intervention [18] to an SIR network model, and some of the closed form network literature (e.g., [19]) may allow a single-world approach. However, our method generalizes this work by explicitly modeling time structure, modeling all state transitions instead of just infection, and allowing additional models and interventions. Auckley et al. explore the relationship between causal diagrams and compartmental models [20], and one could think of our complete graph as extracting unraveling a SWIG from a compartmental model.

Here we focus on a common class of discrete time stochastic state based models to illustrate our method, but the diversity of models used in practice is far greater. While the general idea of the could be applied to any epidemic model, it is the efficient pruning of possible states and events that make this approach tractable. Hence, extension of our approach to certain classes of models (e.g., network models, simple agent base approaches) will be easier than others (e.g., continuous time event driven simulations), and will be for all intents and purposes impossible for some very complex sim-ulation strategies (e‥g, agent based models where disease state changes behaviour). Even within the class of models we consider, for larger meta-population models and other models with a larger number of compartments computational limitations may prevent practical use without significant speedups.

Going forward, there are clear opportunities to improve the applicability and practical utility of this work by optimizing computation and refining pruning assumptions. Also, the current software package is somewhat limiting in the types of control strategies that can be implemented, and we plan to add methods for users to specify their own classes of interventions and appropriate pruning assumptions. The method can also be trivially extended to situations where we are sampling over parameter uncertainty as well as epidemic stochasticity. Perhaps most excitingly, conceptual parallels between our approach and data augmentation approaches (e.g., [21]) suggest ways in which we might more precisely explore how interventions might have worked in real world epidemics.

Precisely characterizing the potential of impacts of interventions is often an important component of scenario modeling analyses for public health, in both pathogen emergent and elimination scenarios. We hope that the methods we develop here lay the groundwork for broadening the set of methods available to achieve this goal, for allowing more precisely targeted inference in a variety of complex scenarios.

